# The transcriptomic architecture of the human cerebral cortex

**DOI:** 10.1101/2024.06.20.599687

**Authors:** Thomas Nickl-Jockschat, Stephen Baek, Zeru J. Peterson, Jerome Charton, Milosch Pavic, Meredith Nastruz, Ethan Rooke, Rachel J. Sharkey, Simon B. Eickhoff, Mike Hawrylycz, Ted Abel

**Author notes:** **Corresponding author** Prof. Dr. med. Thomas Nickl-Jockschat Department of Psychiatry and Psychotherapy Otto-von-Guericke University Leipziger Str. 44 39112 Magdeburg Germany.

## Abstract

For over a century, scientists have been attempting to map the human cerebral cortex, however, they have not taken into account the complex molecular structure of the cortex, which is only beginning to be understood. Here, we parcellate the human cerebral cortex using a machine learning (ML) approach to define its transcriptomic architecture, revealing a multi-resolution organization across individuals. The transcriptomically-derived spatial patterns of gene expression separate the cortex into three major regions, frontal, temporal and parietooccipital, with smaller subregions appearing at lower levels of the transcriptomic hierarchy. The core regions, which remain stable across different hierarchical levels, are physiologically associated with language, emotion regulation, social cognition, motor and visuospatial processing and planning. Importantly, some core regions cross structural and anatomical boundaries identified in previous parcellations of the cortex, revealing that the transcriptomic architecture of the cortex is closely linked to human-specific higher cognitive function.

## Introduction

The human cerebral cortex is by far the most complex biological organ known, harboring higher cognitive functions that are unique for our species and touch to the core of our self-conception as human beings. These unrivaled features have attracted the attention of neuroanatomists for more than a century. Since their start in the era of Korbinian Brodmann^1^ and Santiago Ramón y Cajal^2^, attempts to map the cortex have evolved from early approaches relying on histological criteria and cell morphology^3^ to multi-modal parcellations that capitalize on cutting-edge magnetic resonance imaging (MRI) techniques^4,5^. Modern atlases of the cortex provide a plethora of data about structure and function of the cortex, ranging from three-dimensional cytoarchitectonic maps at a cellular resolution^6–9^ over large functional neuroimaging data bases^10,11^ to advanced simultaneous analyses of multiple different microscopic and imaging modalities^12^. While these maps offer a massive amount of information about the neuronal circuits and cell types composing the cortex, they ignore the transcriptome as major force that shapes the structure and function of these very circuits. Gene expression is a crucial factor for the proper development and functioning of the brain. Experiments in rodents indicate that dynamic changes of transcriptomic patterns are, for example, essential for cognition^13–15^, affective processing^16^, addiction^17^ and the initiation of behaviors^18^. Parcellations based on gene expression patterns, thus, promise unprecedented insights into how the molecular architecture of the brain provides a foundation for cerebral structure and function.

A systematic mapping of spatial transcriptomic patterns, however, has met important obstacles. First, gene expression atlases of the human cortex are largely confined to post-mortem samples, and individual brain donors differ regarding sex, ethnicity, medical history, age at and cause of death, etc. These confounds create a significant amount of unspecific variance between donors that makes it hard to detect common transcriptomic signatures of brain regions across individuals. Second, the comparatively small number of available donor brains significantly adds to the problem of noise, as confounding factors do not average out easily as they would in larger cohorts. Finally, the sheer size of the human cortex and the limitations of contemporary techniques to study complex transcriptomic patterns often require discontinuous mapping, leaving unmapped gaps, for which no data exist.

To address these major obstacles for mapping the transcriptomic architecture of the human cortex, we moved beyond classical univariate statistics to techniques from deep learning and relied upon the Allen Human Brain Atlas^19^ as a data source for brain-wide gene expression. In a first step, we used spectral clustering to identify spatially coherent transcriptomic patterns within the human cortex. We then explored whether there was a distinct transcriptomic hierarchy within those transcriptomic patterns. Finally, we focused on spatially coherent sets of samples that share distinct gene expression patterns across multiple levels of hierarchy, so called core Transcriptomic Brain Regions (core TBRs). Together, this data provides the first data-driven global transcriptomic parcellation of the human cortex.

## Results

### Gene expression patterns form spatially coherent Transcriptomic Brain Regions with a distinct hierarchy

To map transcriptomic patterns across the human cortex, we relied upon data provided by the Allen Human Brain Atlas. The Allen Human Brain Atlas provides a unique resource, as it contains expression levels of >62,000 genes and isoforms from 3,702 samples (punch biopsies) taken from 6 different donor brains. The exact locations, from which the samples were taken, have been mapped to MNI space^20,21^, a standard anatomical reference space in neuroimaging. We used spectral clustering to identify regionally specific transcriptomic patterns that were conserved across donors. For this approach, we treated each sample as a high-dimensional vector, summarizing expression levels of all genes detected at a given anatomic location and the distance of the sample to its neighbors. In order to interpolate gene expression patterns between distinct samples and, thus, allow a continuous mapping, we relied upon the fast marching method^22^ to compute geodesic distances between sample positions along the cortical surface. In contrast to conventional Euclidian approaches, the fast marching method factors in the curvature of the cortex when computing distances between samples. Based on the geodesic distances computed, we then constructed a graph representation of the samples, by connecting neighboring samples with edges. Each graph node was assigned with the gene expression vectors from their corresponding sample, and the edges were weighted with the cosine similarity (L2 norm) of gene expressions between the nodes connected by those edges. This step allowed us to identify spatially coherent sets of samples that shared distinct gene expression patterns (Transcriptomic Brain Regions, TBRs).

To explore a potential hierarchy within the transcriptomic architecture of the human cortex, we performed spectral clustering^23^ with the number of clusters, k, gradually increasing in the range 11 to 101. This approach allowed us to characterize the multiresolution structure of neuroanatomy and its relationship to gene expression in the cortex. At the coarsest level (k=11), our approach yields three major TBRs that covered nearly the entire cortex. Two of these regions are largely identical with the frontal and the temporal lobe, while the third encompassed the occipital and the parietal lobe. By increasing the granularity of our clustering, smaller clusters emerged within these three major TBRs. The large occipito-parietal TBR, for example, decomposes into three regions: one that covers the lateral parts of the occipital and the parietal lobe, one located in the dorsal parietal lobe, and a third that is nearly identical with the primary visual cortex. Of note, the TBR at the occipital pole remains largely unchanged over multiple hierarchical levels. The frontal and the temporal major TBRs show similar hierarchical patterning. In the frontal lobe, we observed a TBR largely identical with the ventromedial prefrontal cortex at the next hierarchical level, while a TBR in the motor cortex emerged only after two more clustering steps. The temporal lobe shows comparatively large TBRs throughout the initial clustering, while more fine grain structures such as the temporal pole, delineate at higher clustering levels. In contrast, in the insular lobe, two TBRs, delineating the anterior and the posterior insula, emerge early and are largely conserved across our analyses (**Figure 1**). We conducted exploratory analyses at k=80 to delineate the level of conservation of TBRs across individual brains and to investigate effects of brain laterality. In the 6 left hemispheres processed for this atlas, we found largely identical TBRs across donors (**Suppl. Figure 1**). For both brains with two processed hemispheres, we found asymmetrical TBRs, mainly reflecting a different parcellation of TBRs in language-related brain regions (**Suppl. Figure 2**).

**Figure 1.**
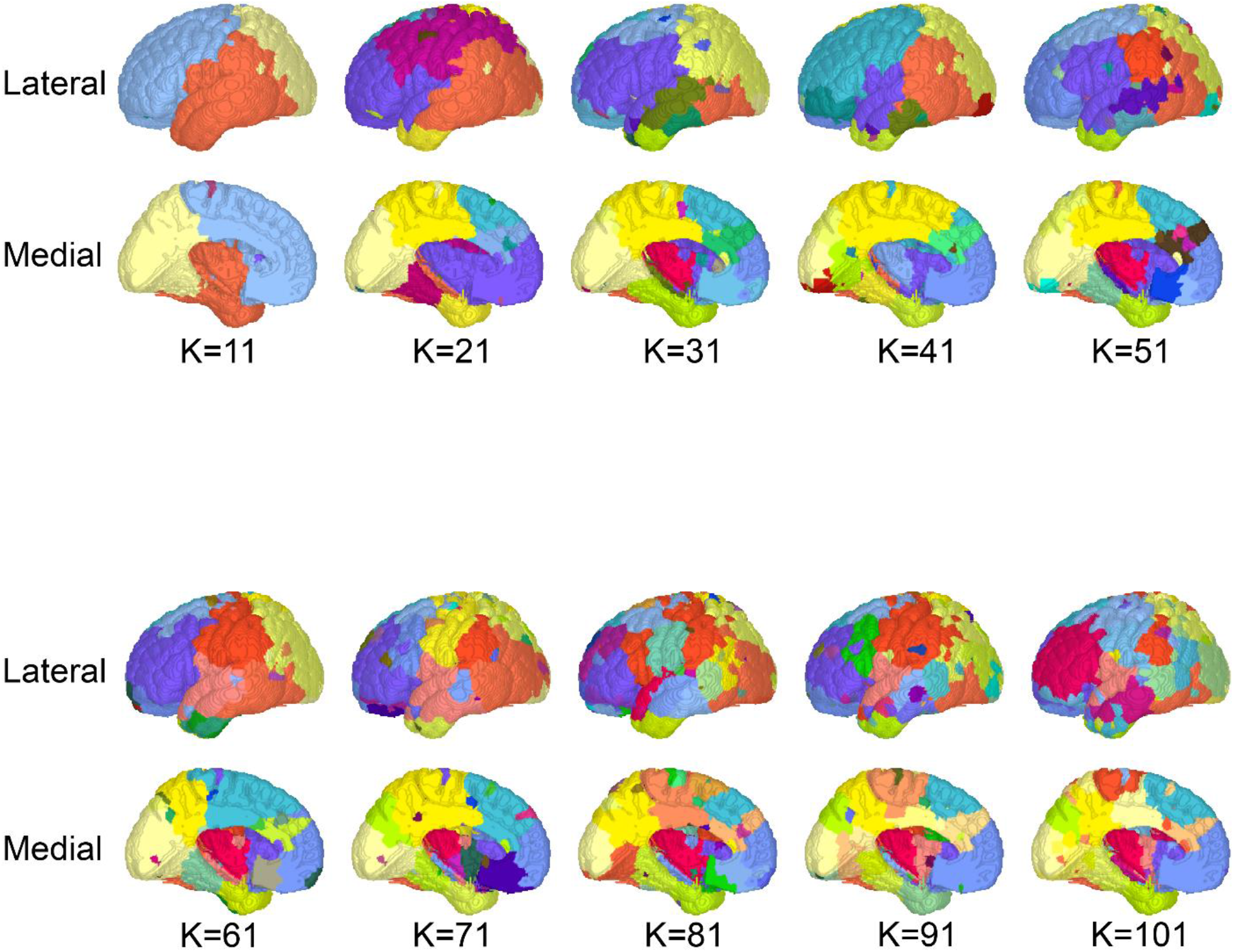
A hierarchical parcellation of the cortex based on transcriptomic patterns across brain donors. Depicted are parcellations of the human cortex from a lateral, medial, ventral and rostral perspective (top to bottom). Parcellations with increasing levels of granularity are shown from left to right, starting with a k=11 to.k=101. At the coarsest level of parcellation, this approach identified three major transcriptomic brain regions (TBRs) that were largely identical with the frontal and the temporal lobe, while one encompassed the parietooccipital cortices. At higher levels of granularity, these clusters split up into smaller filial clusters. Of note, various TBRs remained largely unchanged over different hierarchical levels.

We next investigated whether parcellations at different clustering levels suggested a distinct hierarchy for transcriptomic patterns within the cortex. Our analysis was based upon the hypothesis that a transcriptomic hierarchy would result in larger TBRs decomposing into smaller ones with increasing clustering granularity while retaining overall regional boundaries, and, hence, the combined shape and size of the resulting TBRs at higher granularity would largely recapitulate the parenting TBR. We used the Sørensen-Dice coefficient^24,25^ to determine the overlap between parenting and resulting TBRs. Importantly, this analysis has the advantage that, unlike hierarchical clustering, it does not enforce a hierarchy within the data, but constitutes an approach independently obtained from our initial clustering to validate or falsify our hypothesis. We found evidence for a largely stable transcriptomic hierarchy across most clustering levels. TBRs that did not follow that hierarchical pattern were mainly located at the border between the medial frontal lobe and the insula, and in parieto-occipital regions (**Figure 2**).

**Figure 2.**
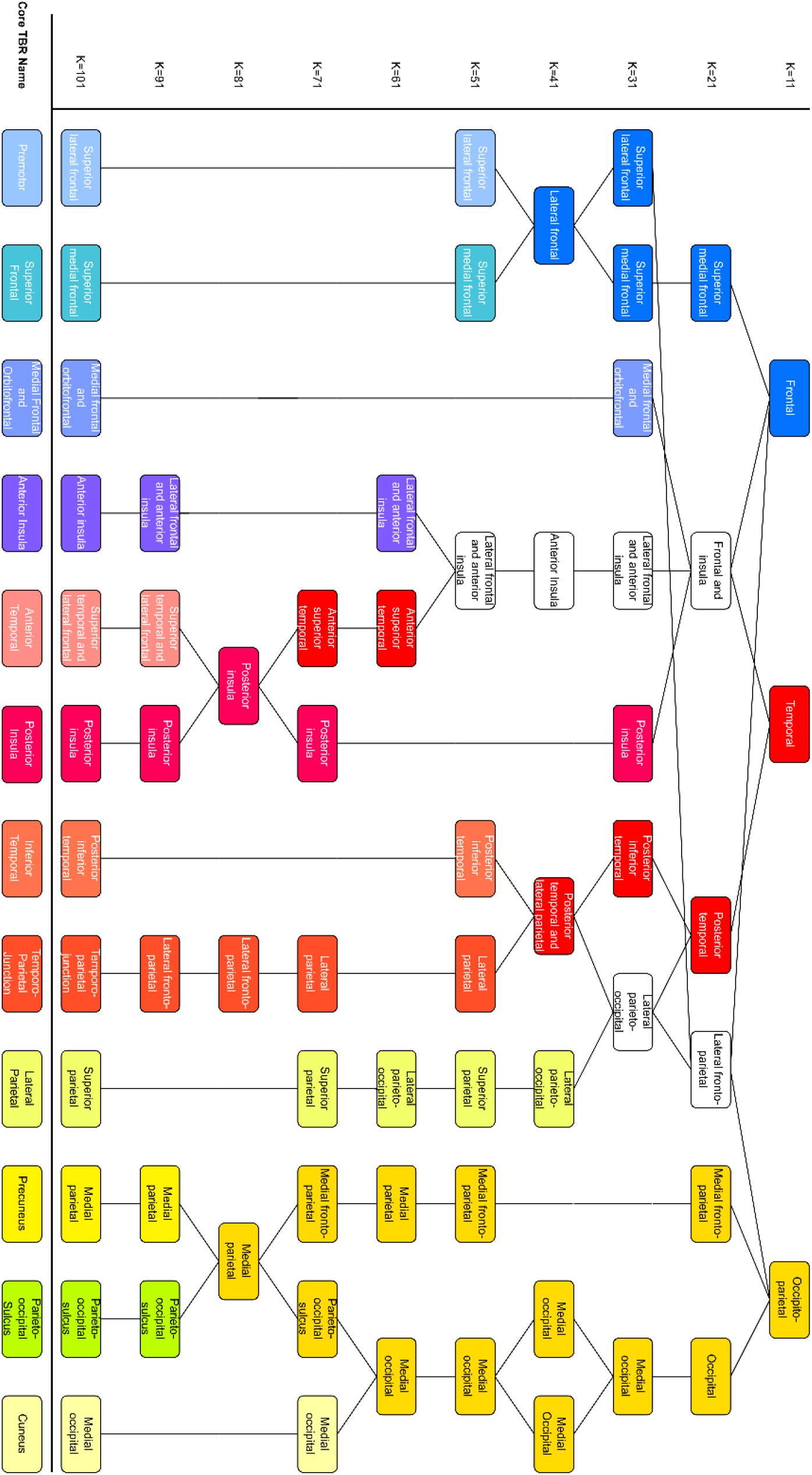
Schematic overview over a hierarchy of spatial transcriptomic patterns in the cortex. Our parcellation of the cortex showed a largely stable transcriptomic hierarchy across most clustering levels. TBRs that did not follow that hierarchical pattern were mainly located at the border between the medial frontal lobe and the insula, and in parieto-occipital regions (Figure 2).

### Transcriptomic Brain Regions that remain stable across different hierarchical levels are associated with higher cognitive functions

Significantly, we found 12 TBRs to remain stable across different hierarchical levels (**Figure 3**). These core TBRs were distributed across the entire cortical surface. Core TBRs were found around the occipital pole, precuneus, parietooccipital sulcus, premotor, superior frontal regions and the ventromedial prefrontal cortex (vmPFC), lateral parietal regions, the anterior temporal lobe, temporoparietal junction, and the anterior and posterior insula. To objectify whether these core TBRs were related to the functional architecture of the brain, we applied functional decoding^11^. This is a data-driven approach relying on the BrainMap functional neuroimaging database^10^, which assigns functional labels to a given brain region based on recorded brain activation patterns and the paradigms that led to these activations in neuroimaging studies. Since this approach only assigns a given physiological function to a brain region, if a related experimental task leads to activations significantly above chance, functional decoding allows inference on how closely TBRs are related to distinct brain functions. Core TBRs were associated with four major physiological domains: language (anterior temporal lobe and posterior insula), emotion regulation (anterior insula, vmPFC, parietooccipital sulcus, temporoparietal junction and superior frontal region), social cognition (vmPFC, precuneus and superior frontal region), motor and visuospatial processing and planning (cuneus, inferior temporal, lateral parietal, precuneus and premotor region). It should be noted that, although linked to brain function, several core TBRs crossed classical structural and anatomical boundaries of the cortex, suggesting a closer link to higher cognitive functions than cellular or classical neuroanatomy.

**Figure 3.**
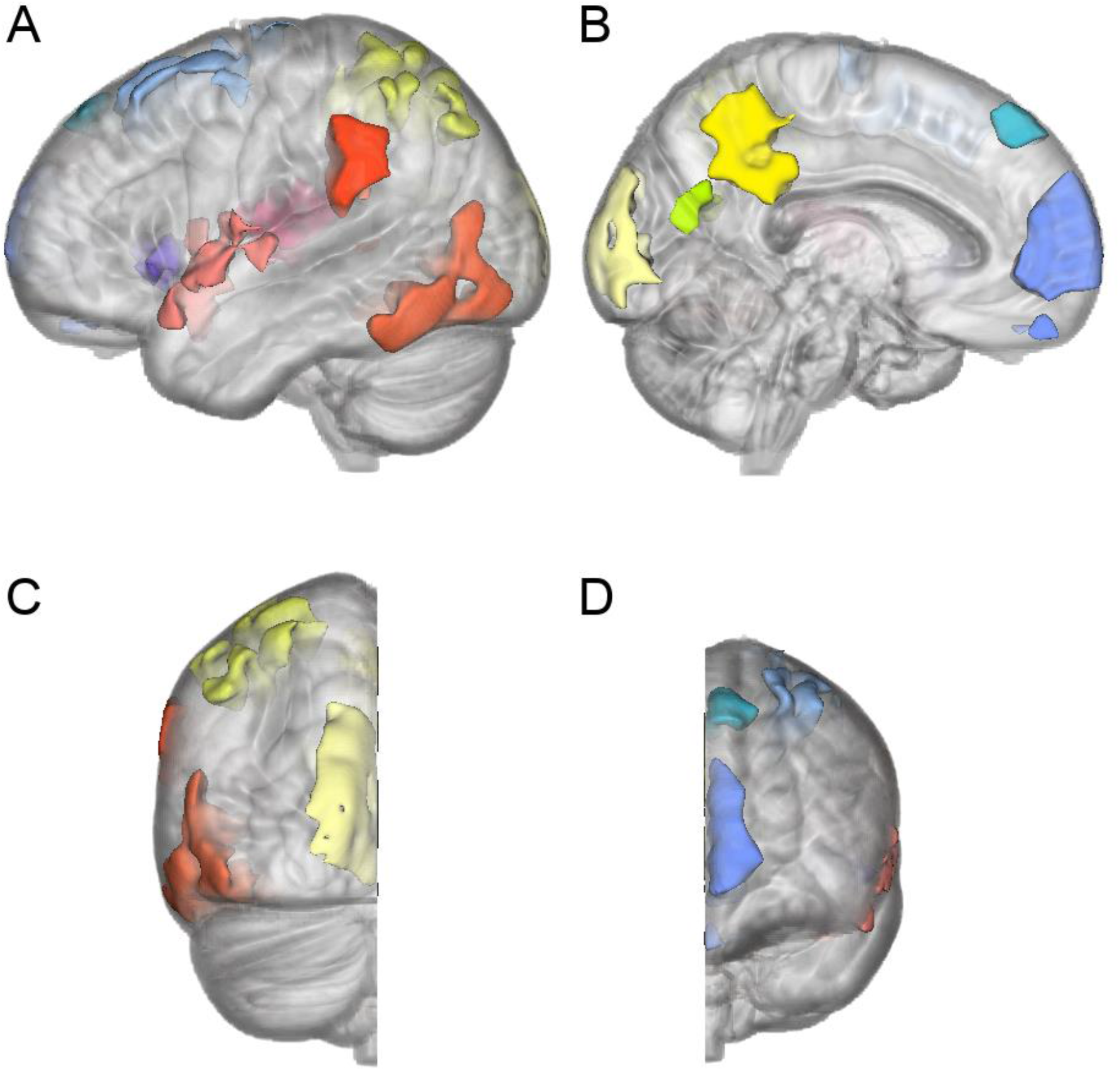
Overview over core transcriptomic brain regions (TBR). Our analyses yielded 12 TBRs that remained stable (= “undivided”) over all hierarchical clustering steps. Depicted are these “core TBRs” from a lateral (A), media (B)l, posterior (C) and frontal (D) perspective. We found core TBRs around the occipital pole, precuneus, parietooccipital sulcus, premotor, superior frontal regions and the ventromedial prefrontal cortex (vmPFC), lateral parietal regions, the anterior temporal lobe, temporoparietal junction, and the anterior and posterior insula. Of note, a data-driven assignment of physiological functions to a given brain region yielded associations with higher cognitive domains: language (anterior temporal lobe and posterior insula), emotion regulation (anterior insula, vmPFC, parietooccipital sulcus, temporoparietal junction and superior frontal region), social cognition (vmPFC, precuneus and superior frontal region), motor and visuospatial processing and planning (cuneus, inferior temporal, lateral parietal, precuneus and premotor region). It should be noted that, although linked to brain function, several core TBRs crossed clasical structural and anatomical boundaries of the cortex, suggesting a closer link to higher cognitive functions than cellular or classical neuroanatomy.

## Discussion

The transcriptomic hierarchy of the mature cortex in adulthood might reflect differentiation processes during neurodevelopment. The three main TBRs at the coarsest level of parcellation (k=11) – frontal, temporal and occipitoparietal – parallel spatial expression patterns of three key morphogene families during the earliest stage of cortical development. Wnts/BMPs, for example, show a predominantly occipitoparietal expression pattern^26–29^, while Shh exhibits temporal^30–32^ and FGF8 frontal expression^33–35^, respectively. Further research relating morphogene expression during neurodevelopment to the mature transcriptomic architecture should help to gain a better understanding of how the molecular basis of cortical structure and function emerges.

Gene expression is reflective of distinct cell types and, hence, the cellular composition of a given brain region will influence regional transcriptomic patterns^36–38^. Neuronal and glial activity, on the other hand, e.g., due to cognitive and affective processes or the initiation of behaviors, have been identified as major modulators of gene expression in animal models^39,40^. The close association of our core TBRs with functional, rather than known macro- or microstructural, parcellations of the brain, gene expression appears mainly driven by functional processes. This influence of brain function on transcriptomic profiles is sufficiently pronounced that it drives regional similarity even in post-mortem samples.

Of note, core TBRs demonstrating stability across multiple clustering levels were mainly related to higher cognitive functions, such as language, emotion regulation, and social cognition. This was contrary to our initial expectations, as we hypothesized that TBRs associated with primary sensory and motor regions would be among the most conserved. However, our findings highlight the obvious physiological importance of these higher brain functions. Language, social cognition, and emotion regulation are key processes that enable humans to form relationships within larger groups and are, therefore, a major requirement for the emergence of culture^41–43^. Social cooperation within more and more complex societies has proven to be a major evolutionary advantage in humans^44^. Our findings of highly conserved transcriptomic patterns related to social processes could provide the molecular foundation of neural processes that enable humans these complex social interactions.

In summary, investigating the structure of the adult human brain based on transcriptomic derived brain parcellation demonstrates evidence for a transcriptomic hierarchy conserved across different donor brains. While recent scientific approaches have focused on extracting cortical parcellations from neuroimaging results, our work provides a wholly new angle and demonstrates a brain parcellation that is based on transcriptomic patterns which are conserved across individuals. Given the striking heterogeneity of donors regarding age at and cause of death, ethnic background, and sex, a conserved parcellation appears as non-trivial. These results will provide the foundation for future studies exploring the transcriptomic architecture on a cell-specific level such as data derived through the recently launched BRAIN Initiative Cell Atlas Network (BICAN) to profile cell types in the human and non-human primate.

## Methods

### Data sources

For our method we used data from 6 post-mortem brains collected by the Allen Institute for Brain Science. Since whole brain data existed for only two of the brains, we instead focused our analysis on the left hemispheres, which were collected for all 6 donors. For each of these brains, microarray punches (samples) were collected across the cortex and the location recorded in MNI space, with a total of 3,702 total samples collected across the whole brain, and 1,111 samples within the grey matter of the left hemispheres of all 6 donors. Each of these microarray samples contained the transcriptomic data for 58,692 genes. More detailed methods of collection used for the data in this analysis is described in more detail in Hawrylycz et al., 2012^19^, and Hawrylycz et al., 2015^45^.

### Fast marching calculation of distance between cortical samples

For our data, each sample constituted a node in the graph at a location defined in MNI space, and edges were defined using the 12 nearest neighbors to each sample based on geodesic distance. The first step of extracting regional transcriptomic patterns was to isolate which samples were located within the cortex. To do this we used a gray matter mask from the MNI brain atlas^20,21^ to exclude non-cortical samples from the sample list, leaving a total of 1,111 samples within the cortex out of the original 3,702 samples. We then used the gray matter mask as our boundary for measuring geodesic distance between cortical samples using a fast marching algorithm^22^. This allowed us to calculate geodesic distance between samples along the cortical gyri, rather than calculating Euclidean distance which might cross through anatomical boundaries such as the sulci between cortical gyri. A total of 12 nearest neighbors were calculated per sample using this method, and these nearest neighbors were then used to generate the edges between samples and calculate the cosine similarity degree matrix for the spectral clustering.

### Spectral embedding of transcriptomic data

After generating the list of nearest neighbors in the cortex, we calculated the cosine similarity (Eq. 1) for each sample to its neighbors, where Ai is the vector of transcriptomic data for the sample of interest, and Bi is the vector of transcriptomic data from a neighboring sample. The entire vector of transcriptomic information as the input for the cosine similarity calculation. We created a sparse weighted adjacency matrix using all 1,111 cortical samples, populated with data from their 12 nearest neighbors as well as a degree matrix showing how many connections each sample makes to other cortical samples. A Laplacian matrix was generated by subtracting the adjacency matrix from the degree matrix, min-max normalized, and used to calculate the eigenvectors and eigenvalues for the graph.

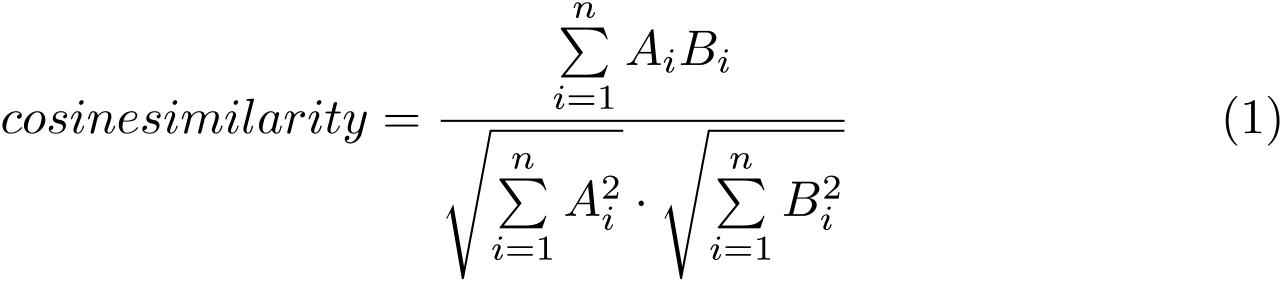

### Spectral clustering of embedded data

To perform the clustering of our data we selected the 300 largest magnitude eigenvalues and ran their corresponding eigenvectors through a *k*-means clustering algorithm, sequentially increasing the *k* by 10 between *k*=11 and *k*=101 to look for hierarchical patterns in the clustering. Results of the clustering were then mapped back into MNI space using the original coordinates for each clustered sample, and a label mask was generated where each region is labeled based on the clustering. We then took each of the label masks from *k*=11 to *k*=101 and added the masks together to look for overlapping regions. By finding the samples that were consistently clustered together across various levels of clustering, we were able to generate Transcriptomic Brain Regions (TBRs) defined by their conserved transcriptomic patterns within a region.

### Functional characterization of TBRs

To further characterize the brain functions of core TBRs, we analyzed the associated BrainMap meta-data^10,11^ (http://www.brainmap.org) for significant associations. The key idea behind this approach is to identify all experiments that activate a particular region of interest – in this case a core TBR - and then analyze the experimental meta-data describing the experimental settings that were employed in these^46–48^. This allows statistical inference on the type of tasks that evoke activation in a particular region, allowing the data-driven identification of brain-behavior relationships and avoiding the problems that come with functional assignments to brain regions that are based on subjective hypothesis.

In this study, we used behavioral domains (BD) from the BrainMap database that describe the cognitive processes probed by an experiment. In the forward inference approach, the functional profile was determined by identifying taxonomic labels for which the probability of finding activation in the respective region/set of regions was significantly higher than the overall (*a priori*) chance across the entire database. That is, we tested whether the conditional probability of activation given a particular label [P(Activation|Task)] was higher than the baseline probability of activating the region(s) in question *per se* [P(Activation)]. Significance was established using a binomial test [*p* < 0.05, corrected for multiple comparisons using false discovery rate (FDR)]. In the reverse inference approach, the functional profile was determined by identifying the most likely behavioral domains, given activation in a particular region/set of regions. This likelihood P(Task|Activation) can be derived from P(Activation|Task) as well as P(Task) and P(Activation) using Bayes’ rule. Significance (at *p* < 0.05, corrected for multiple comparisons using FDR) was then assessed by means of a chi-squared test.

### Author contributions

Conceptualization: T.N.-J., S.B., T.A.

Methodology: S.B., Z.J.P., J.C.

Investigation: T.N,-J., S.B., Z.J.P., J.C., M.P., M.N., E.R., R.J.S.

Visualization: J.C., Z.J.P., R.J.S.

Supervision: T.N.-J., S.B., T.A:

Data interpretation: T.N.-J., S.B.E., M.H., T.A.

Writing – original draft: T.N.J., T.A

## Acknowlegdements

Thank you to A. Bayly for assistance with colour selection for the figures.

## Data and materials availability

All data needed to evaluate the conclusions in the paper are present in the paper, The original data from the Allen Human Brain Atlas can be found at

